# Local neuronal excitation and global inhibition during epileptic fast ripples in humans

**DOI:** 10.1101/2021.09.09.459695

**Authors:** Jonathan Curot, Emmanuel Barbeau, Elodie Despouy, Marie Denuelle, Jean Christophe Sol, Jean-Albert Lotterie, Luc Valton, Adrien Peyrache

## Abstract

Understanding the neuronal basis of epileptiform activity is a major challenge in neurology. Interictal epileptiform discharges are associated with fast ripples (FRs, >200 Hz) in the local field potential (LFP) and are a promising marker of the epileptogenic zone. Here, by using a novel hybrid macro-micro depth electrode, combining classic depth recording of LFP and two or three tetrodes enabling up to 15 neurons in local circuits to be recorded simultaneously, we have characterized neuronal responses to FRs on the same hybrid and other electrodes targeting other brain regions. While FRs were associated with increased neuronal activity in local circuits only, they were followed by inhibition in large-scale networks. Neuronal responses to FRs were homogeneous in local networks but differed across brain areas. Similarly, post-FR inhibition varied across recording locations and subjects and was shorter than typical inter-FR intervals, suggesting that this inhibition is a fundamental refractory process for the networks. These findings demonstrate that FRs engage local and global networks and point to network features that pave the way for new diagnostic and therapeutic strategies.

## Introduction

Epilepsy is a major chronic neurological condition, which can arise from genetic mutations, developmental malformations, or acquired cortical lesions from heterogeneous causes (e.g. traumatic, infectious, or vascular events). Epileptic patients suffering from a drug-resistant form of epilepsy may necessitate surgical intervention to stop, or at least diminish, the occurrence of seizures ^1^. When it is believed that there is a specific epileptogenic zone (EZ) that can be removed by surgery ^2^, a non-invasive and sometimes invasive workup – depending on the complexity of each case – is performed to determine where and how seizures begin ^3^, including stereoelectroencephalography (SEEG).

While the recording of spontaneous seizures remains the gold standard to define the EZ, this procedure is time-consuming, logistically demanding, and uncomfortable for the patients. These limitations could potentially be overcome by analyzing the interictal periods. Fast-ripples (FRs, 200 Hz-600 Hz) are often associated with interictal epileptic discharges (IEDs, i.e. “epileptic spikes” or “sharp waves”) and are promising candidates for an interictal biomarker of the EZ ^4–9^.

FRs are generally considered to be more specific and more focal than IEDs alone, which can be found beyond the EZ ^8,10,11^ and are observed in the local field potential (LFP) over several millimeters ^12–14^. In support of this view, resecting areas with the highest rate of FRs has been linked to a better surgical outcome ^6,15–18^. The specificity of FRs in identifying the EZ was recently questioned ^19,20^. However, these conclusions were drawn from observations of FRs on macroelectrodes. Overall, the study of FRs on micro-electrodes remains limited ^4,21–25^. It has thus become crucial to not only further characterize FRs at the micro-circuit level but to unravel the neuronal dynamics underlying these events and their consequences for neuronal activity in other cortical areas.

Animal work has provided a wealth of data regarding the generation of FRs and hippocampal ripples (100-200Hz), which naturally occur in healthy tissue ^26–32^. This was achieved in part with techniques such as multi-channel electrophysiology, which allows for monitoring the activity of populations of neurons. In humans, neuronal recordings are achieved in deep structures with single microwires, offering only limited separation power of neuronal waveforms, or on the cortical surface with electrode arrays that are usually distant from the EZ ^33^. While previous work on the relationship between IEDs and neuronal activity reported highly diverse responses ^34,35^, the specificity of local and long-range population responses of FRs remains unclear. Here, we used a new type of intracranial electrode equipped with microwires bundled in tetrode configurations ^36^. Tetrodes (composed of four microwires bundled together) lead to a high yield in the number of isolated units ^37^ and provide a much higher spatial resolution to determine the source localization of FR generators. As these microwires are guided through regular intracerebral depth electrodes routinely used for the localization of the EZ, FRs can be recorded at both micro and macro scales simultaneously, with a high spatial correspondence, in relation to neuronal activity.

Interictal FRs were detected in subjects implanted with these new “hybrid” electrodes. We first demonstrate that FRs are associated with strong neuronal excitability in the EZ. Next, we show that neurons outside the EZ are not excited by the FRs, yet they decrease their firing after FRs. Last, we show that post-FR firing decreases can last up to one second and are related to the inter-FR intervals. We suggest that this global post-FR inhibition is part of a refractory mechanism preventing FRs from developing into more general seizures.

## Results

Drug-refractory epileptic patients (n = 9) were implanted with 8-15 depth electrodes to accurately localize the EZ. Following implantation, SEEG recordings were performed for 5 to 12 days. Our analysis epochs were extracted from the five first days. The location of implantation was determined according to the clinical hypothesis of EZ location based on the non-invasive workup.

### A new macro-micro electrode to simultaneously record local field potentials (LFP) and multiple single units in deep local brain circuits

Three to four of the implanted depth electrodes were a hybrid version combining macro-contacts and 2-3 tetrodes between the two deepest macro-contacts (8-12 tetrodes per subject; see Methods section). The interictal signal was recorded for one hour at a high sampling rate (30 kHz) in each subject during the day-time period. FRs (200-600 Hz) were visually detected throughout this one-hour signal ^8,38^.

We selected 9 subjects (clinical details in the supplementary material) from our entire database who combined a high signal-to-noise ratio, detection of at least 1 FR, and/or detection of at least 1 isolated single unit on one tetrode. Tetrode locations were confirmed by co-registration of preoperative stereotactic MRI and post-operative CT scans (**Fig. 1a**) and the dataset was composed of recordings in the anterior hippocampus (9 hybrids, 21 tetrodes), posterior hippocampus (6, 15), amygdala (8, 19), rhinal cortex (8, 17), temporo-occipital or lingual gyrus (6, 12), temporal pole (3, 6), and superior temporal gyrus (2, 4). Thirty-seven tetrodes were located in the right hemisphere, 49 in the left hemisphere. Both FRs and single units were recorded on 16 tetrodes, FRs only on 31 tetrodes, and action potentials only (without any FRs) on 12 tetrodes (**Fig. 1b**). In total, 2233 FRs were detected in the 9 subjects (4-470 FRs per hybrid).

**Fig. 1.**
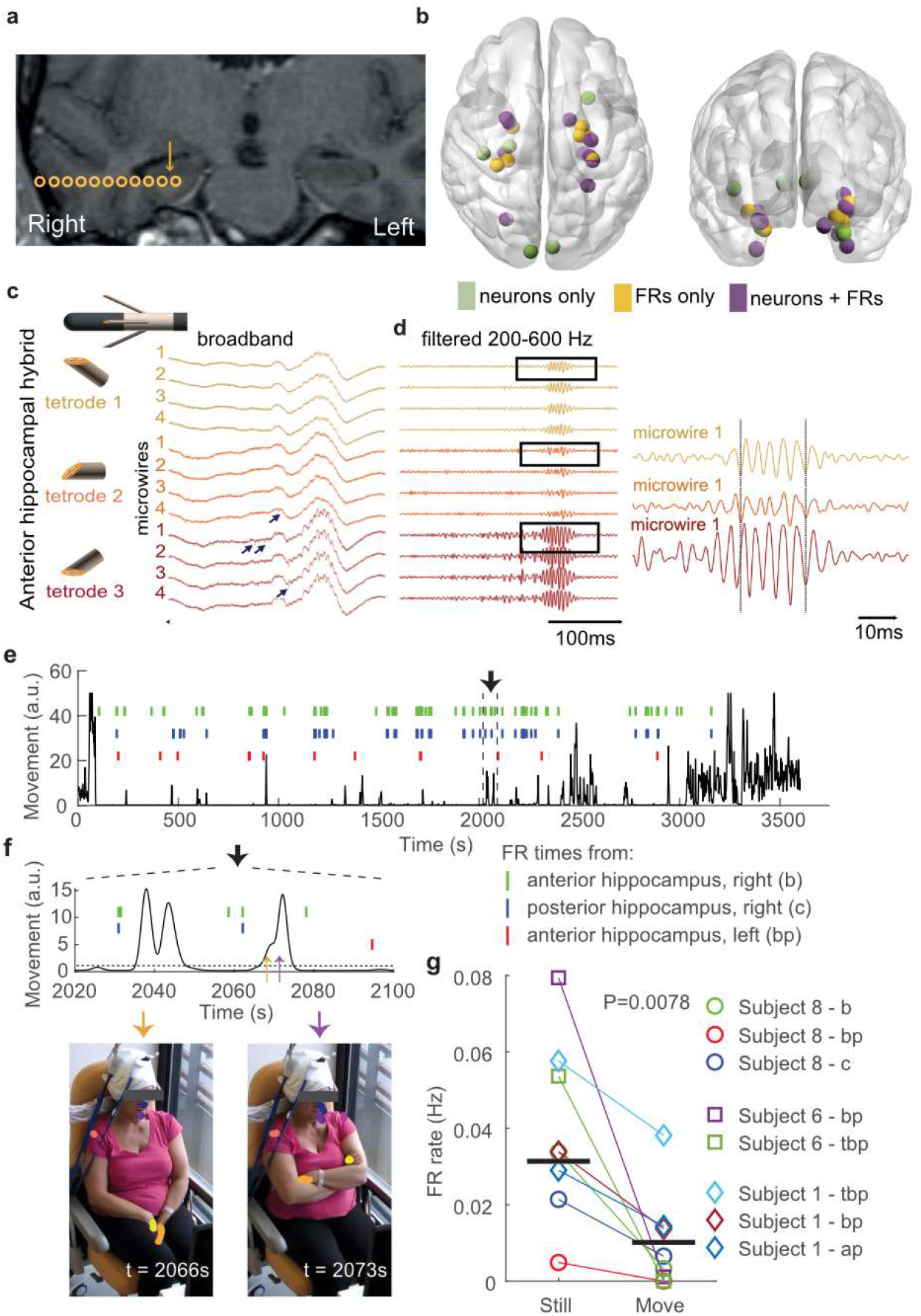
Recording of intracerebral signals with micro-wires at a high sampling rate reveals local FRs and action potentials. **a**. Example of hybrid electrode location in one subject (*Subject 8*). Coronal co-registration MRI CT scan (preoperative 3D T1 MRI with gadolinium and post-operative CT scan). Yellow circles are macro-contacts; the arrow specifies the putative location of tetrodes between the two deepest macro-contacts of the hybrid electrode. **b**. Locations of every tetrode included in the study, referenced in the MNI space. Colors indicate whether FRs, single units (neurons), or both were detected. See Supplementary Table 1 for complete list of locations. **c**. Schematic representation of the deepest macro-contacts and the three tetrodes and example LFP traces (broadband) from the three tetrodes of a hybrid electrode targeting the right anterior hippocampus (*Subject 8*). The IED (positive deflection) is associated with a FR. Note that both IED and FR amplitude are higher on the 3^rd^ tetrode. Black arrows indicate action potentials. **d**. *Left*, same signals as in **c** filtered in the FR frequency band (200-600 Hz). Note that FRs have the same morphological features on the four microwires of each tetrode. *Right*, zoomed trace of the first microwire of each tetrode. Although FRs occur at the same time, moment-to-moment phase difference varies. **e**. Subject upper body movement (see Methods section) and occurrence times of FRs on three hybrid electrodes during a one-hour long recording. **f**. *Top*, temporal zoom of the traces in **e** (delimited by vertical dotted lines). *Bottom*, two video frames extracted at times indicated by the orange and purple arrows in top panel. Note the markers on the head, shoulders, and hands. **g**. FR occurrence rate during stillness and movement for all three subjects with video tracking. Horizontal bars indicate median values.

Importantly, all FRs were associated with an IED (see, for example, **Fig. 1c-d**), which demonstrates that they were not simply physiological ripples. FRs were generally very similar on all four wires of a tetrode. However, FRs often had different amplitudes and were not necessarily in phase on the different tetrodes from the same hybrid (**Fig. 1d**). Because the tetrode tips of a hybrid electrode are separated by at most 2 mm ^36^, these findings suggest that the underlying neuronal generators of FRs are characterized by an anatomical scale in the millimeter range.

Further evidence that FR generators are anatomically small comes from the observation that while FRs detected on the deepest macro-contacts were always associated with FRs on the tetrodes, the opposite was not always true: 82% (range: 38-100%) of FRs detected on tetrodes were associated with detectable FR on the nearest macro-contact (n = 6 hybrid electrodes in three subjects with synchronized macro-contact and tetrode recordings).

The occurrence rate of FRs recorded on clinical macroelectrodes depends on vigilance states, showing the highest incidence during sleep ^18^. Here, we asked whether FRs detected on micro-wires were also modulated by vigilance states at a fine temporal resolution. To this end, we recorded subjects with a camera whose video stream was synchronized with the tetrode electrophysiological signals in three subjects with a high FR incidence rate on at least two hybrid electrodes (see Methods section). While the recordings only included the interictal day-time period in the morning, these three subjects all show a highly variable level of attention during the recording, from fully awake to asleep. We determined their vigilance states by tracking their movements using DeepLabCut ^39^ (see Methods section) (**Fig. 1e,f**). We observed a 3-fold increase of the FR occurrence rate during the immobility period (p = 0.0078, n = 6 hybrid electrodes, Wilcoxon sign rank test; **Fig. 1g**). These findings show that brain states corresponding to – even transient – motor activity are associated with a decrease in the excitability of FR generators.

### Recording FRs and single units in local and global circuits

Raw electrophysiological signals were processed to semi-automatically extract action potentials (**Fig. 2;** see Methods sections). A total of 139 single units were isolated in 9 subjects, corresponding to 17.4 (± 9.1 SD) single units per subject, 7.3 (± 5.1) per hybrid electrode, and 5 (± 3.6) per tetrode (for all subjects, hybrid electrodes, or tetrodes where at least one single unit was detected).

**Fig. 2.**
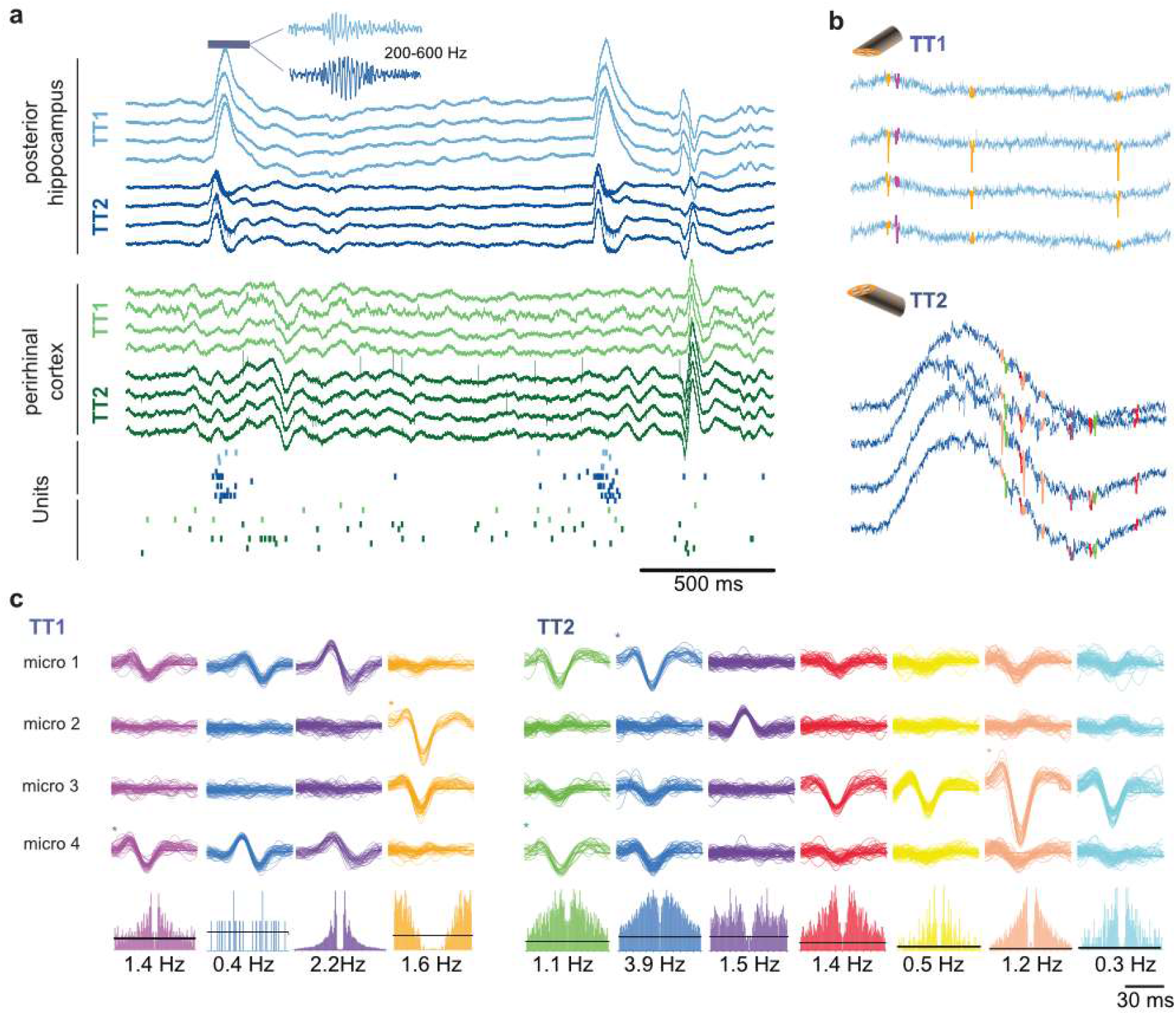
FR and spikes in local and global circuits. **a**. LFP recordings (unfiltered broadband signal) on two hybrid electrodes in *Subject 2*. Two IEDs are visible in the posterior hippocampus, one in the perirhinal cortex. IEDs were associated with FRs, as shown in the inset (200-600 Hz filtered LFP trace). *Bottom*, raster plot of neuronal activity. Each row corresponds to one unit and each dot to one action potential. There is an increase in firing in most units during the IED and FR recording on TT1 and TT2 located in the posterior hippocampus (blue and purple dots). **b**. Action potentials recorded on the posterior hippocampus on TT1 and TT2 immediately before the highlighted FR in **a** (TT1) and during the FR (TT2). Note the FR in the broadband trace. **c**. Sample waveforms of action potentials recorded on TT1 and TT2 in the posterior hippocampus, sorted into putative single units. *Bottom*, auto-correlogram (± 30ms) and mean firing rate of each single unit.

To investigate the relationship between neuronal activity and FRs, we divided the dataset into *local* and *global* interactions, corresponding to neuronal activity in relation to FRs recorded on the same hybrid or another hybrid electrode (n = 78 and 190 single units studied for local and global interaction, respectively, in nine subjects; note that the same neuron can be counted twice if FRs were detected on more than one hybrid).

### Local FR-related excitation but global post-FR inhibition

We next asked how FRs modulated local and global neuronal activity in epileptic subjects. To this end, we computed spike train cross-correlograms in relation to FR occurrence times (that is, average firing rate as a function of time-lag from FRs). Firing rates are broadly distributed in the cortex ^40^, making the direct comparison of cross-correlograms challenging. We therefore normalized cross-correlograms by number of standard deviations from baseline (i.e. average firing rate expected by chance) for each time-lag ^41^ (**Fig. 3a**; see also Methods section and **Supplementary Fig. 1**).

**Fig. 3.**
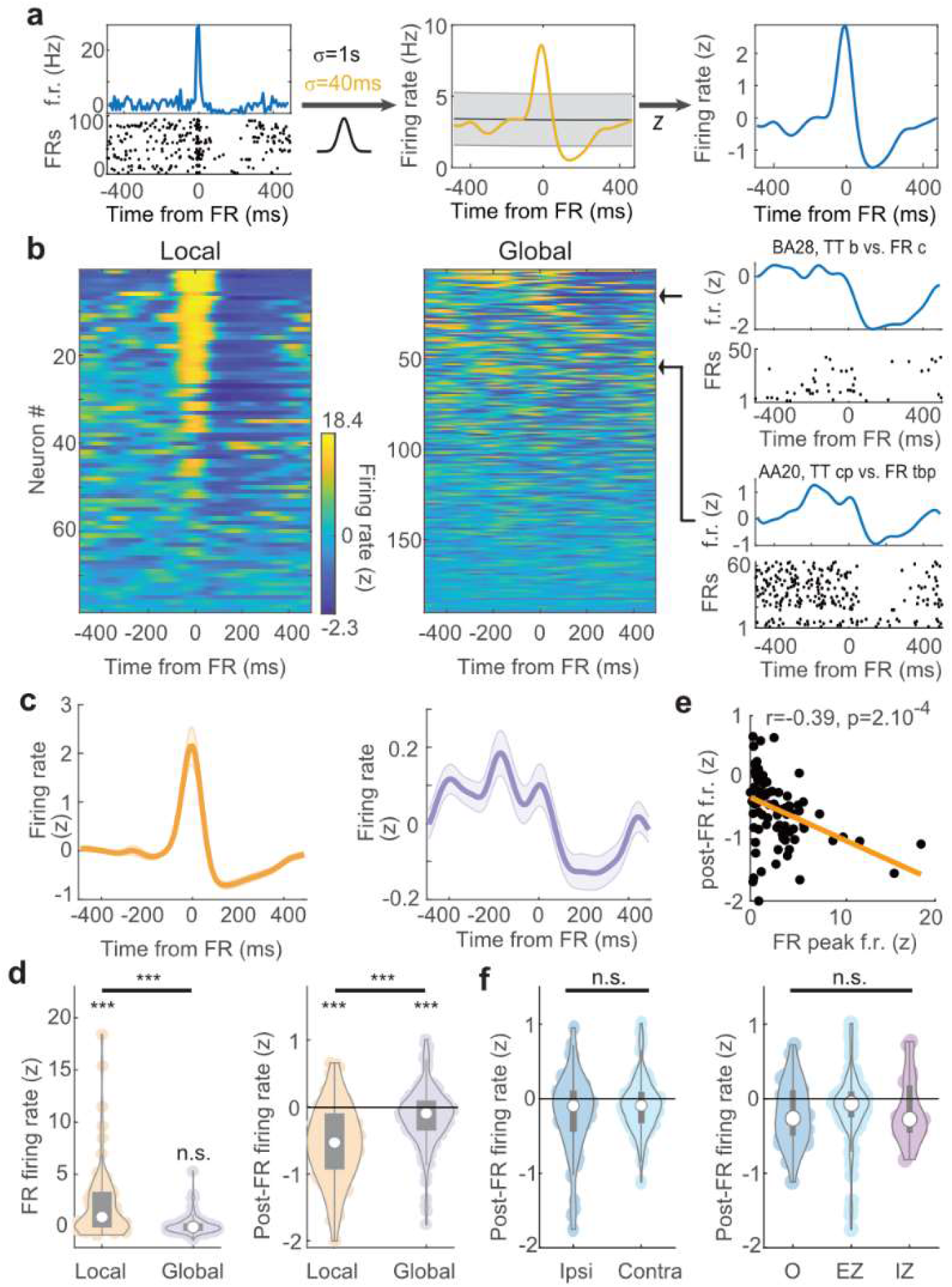
Local and global modulation of neuronal activity with FRs. **a**. Normalization of a sample single unit cross-correlograms relative to time of FRs. Time 0 corresponds to the occurrence of FRs. *Left*, mean firing rate (f.r.) of the neuron relative to FR occurrence time (*top*) and raster plot of FR-by-FR spiking activity (*bottom*); *middle*, the cross-correlogram is convolved with two Gaussian filters of different width; *right*, the z-scored normalized firing rate (see Methods section). **b**. Normalized cross-correlograms of each single unit recorded at the local (same hybrid as FR) and global scale (different hybrid) relative to FRs. Colors display z-scored firing rates. Cross-correlograms are sorted by their overall variance, from maximum (*top*) to minimum (*bottom*). *Right*, two sample single units at the global range (titles indicate location of hybrids from which neuron and FRs were recorded). **c**. Average cross-correlograms at the local (*left*) and the global (*right*) scale. **d**. *Left*, FR peak firing rates at the local and global scale; *right*, same for post-FR firing rates. **e**. Post-FR firing rates as a function of FR peak firing rates in local networks. **f**. *Left*, post-FR firing rates at the global scale for ipsi- and contralaterally located hybrid electrodes (i.e. hybrids on which FRs and neurons were monitored). *Right*, same for the different status of the neuron-recording hybrid electrodes (O: healthy tissue, EZ: epileptogenic zone; IZ: irritative zone). In panels **d** and **f**, white circles indicate the median abd gray rectangles the distribution of the first two quartiles around the median.

At the time of FRs, local neurons, but not neurons from remote hybrid electrodes, showed strong activation (**Fig. 3b-d**; p = 6.6 × 10^−8^, 0.81; n = 78, 190 for local and global interaction, respectively; Wilcoxon signed rank test; local firing greater than global firing: p = 6.7 × 10^−9^, Mann-Whitney U test). In contrast, both the local and global networks showed a decrease in firing rates after FRs (**Fig. 3b-d**; p = 5 × 10^−10^, 2.2 × 10^−5^; Wilcoxon signed rank test). However, the decrease in local firing rate was significantly stronger than for remote neurons (p = 1.3 × 10^−9^, Mann-Whitney U test). These observations provide additional evidence that FR generators are restricted to local circuits. Interestingly, they point to a potential mechanism to dampen activity following FRs in large-scale networks.

We then sought to determine whether the variability in neuronal responses, both the activation of local neurons at times of FRs and the local and global inhibition, could be explained by anatomical and subject-specific factors. We first examined local network responses. A majority of local neurons showed increased firing rates at the time of FRs (**Fig. 3c,d**; 57/78 neurons with z > 0; p = 2 × 10^−5^; binomial test) and post-FR decrease in firing rates (64/78 neurons with z < 0; p = 3 × 10^−9^; binomial test). The FR increase and post-FR decrease of firing rates were correlated (**Fig. 3e**; r = 0.39; p = 2 × 10^−4^; Pearson’s correlation) and, accordingly, local neurons that were activated with the FRs were virtually all followed by a period of decreased firing (51/57 neurons, p = 3 × 10^−10^; binomial test).

Next, we examined two potential sources of variability at the global level: post-FR firing rates depended neither on whether FRs were ipsi- or contra-lateral (p = 0.3; Kruskal-Wallis test) nor on the status of the network (EZ, irritative zone, or healthy cortex; see Methods section) from which neurons were recorded (**Fig. 3f**; p = 0.08; Kruskal-Wallis test), suggesting a global phenomenon that cannot be accounted for by a simple relationship between the FR-generating tissue and the target areas.

### Local homogeneity but global heterogeneity in neuronal responses to FRs

Local networks are strongly activated at the time of FRs, but are all neurons similarly modulated? To address this question, we focused our analysis on three subjects in which at least one hybrid showing FRs had two tetrodes, each with at least three neurons (**Fig. 4a**; total of 10 tetrodes on 6 hybrid electrodes). Neuronal activation at the time of FRs was more similar within each tetrode than across tetrodes (**Fig. 4b**; p = 1.4 × 10^−4^; n = 82 neurons from n = 4, 3, 3 tetrodes from the same hybrids in three different subjects; Kruskal-Wallis test).

**Fig. 4.**
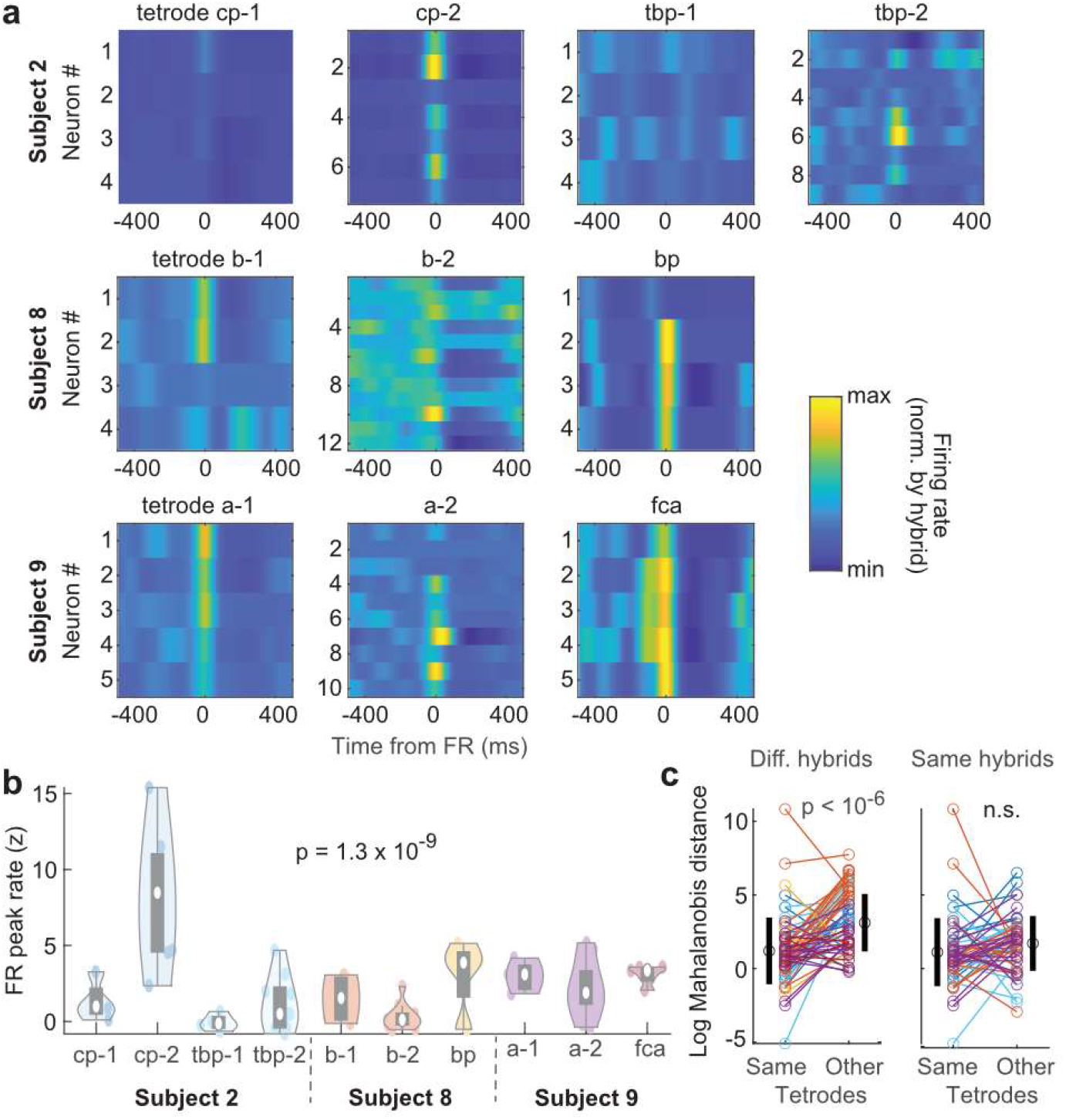
Neuronal modulation by FRs in local networks. **a**. Tetrode-by-tetrode neuronal cross-correlograms in three subjects (total of six hybrid electrodes and 60 single units). Color indicates z-scored firing rates from minimum (blue) to maximum (yellow) on each hybrid electrode. **b**. Distribution of tetrode-by-tetrode peak FR firing rates. White circles indicate the median and gray rectangles the distribution of the first two quartiles around the median. **c**. Comparison of cross-correlograms profiles (i.e. after compressing extreme values) between neurons of the same tetrodes or other tetrodes (see Methods section). *Left*, Log Mahalanobis distance of each neuron to all other neurons from the same tetrodes and average distance to tetrodes on other hybrid electrodes. Black circles and vertical lines on the sides show average and SD, respectively. *Right*, Same as *left* but within tetrode distance in comparison to average distance to tetrodes from the same hybrid. Color of each neuron indicates hybrid electrode of origin, as in **b**.

Although absolute activation visibly differs between tetrodes from the same hybrids (but generally fails to reach statistical significance because of limited sampling), the relative temporal profile of neuronal activity around FRs may still be specific to local networks. To test for this possibility, we first renormalized cross-correlograms to compress the range of extreme values. We then computed the similarity (i.e. Mahalanobis distance, see Methods section) of each neuron to subsets of other neurons from the same or other tetrodes. In each subject, the response profile of each neuron was more similar to the other neurons of the same tetrodes than to the neurons of tetrodes of other hybrid electrodes (**Fig. 4c**; p < 10^− 6^, n = 64, paired t-test), but was similar to neuronal responses on tetrodes from the same hybrid electrodes (p > 0.05, n = 55, paired t-test; **Fig. 4c**). Overall, these observations provide further evidence for the small scale (∼1 mm) of FR generators and suggest that neurons of a given brain area show similar dynamics although absolute activation can differ.

### Circuit-specific post-FR recovery of neuronal activity and FRs

Post-FR inhibition is particularly strong in FR-generating networks, bearing strong similarities with inhibitory currents governing hippocampal activity following physiological ripples ^27,29^ and believed to impose a refractory period on ripple generation. We thus asked whether the post-FR decrease in firing rate was related to the dynamics of FR generation. First, we characterized post-FR inhibition for each single cell by fitting a sigmoid on spike train cross-correlograms relative to FRs (**Fig. 5a**). Post-FR neuronal recovery time was defined as the time of maximum sigmoid slope after the FR (that is, the time at which neurons reached 50% of their baseline firing rate). Only units for which the sigmoid fit converged and for which the recovery time was within the range 50-1500ms were included in the analysis (n = 55 single units).

**Fig. 5.**
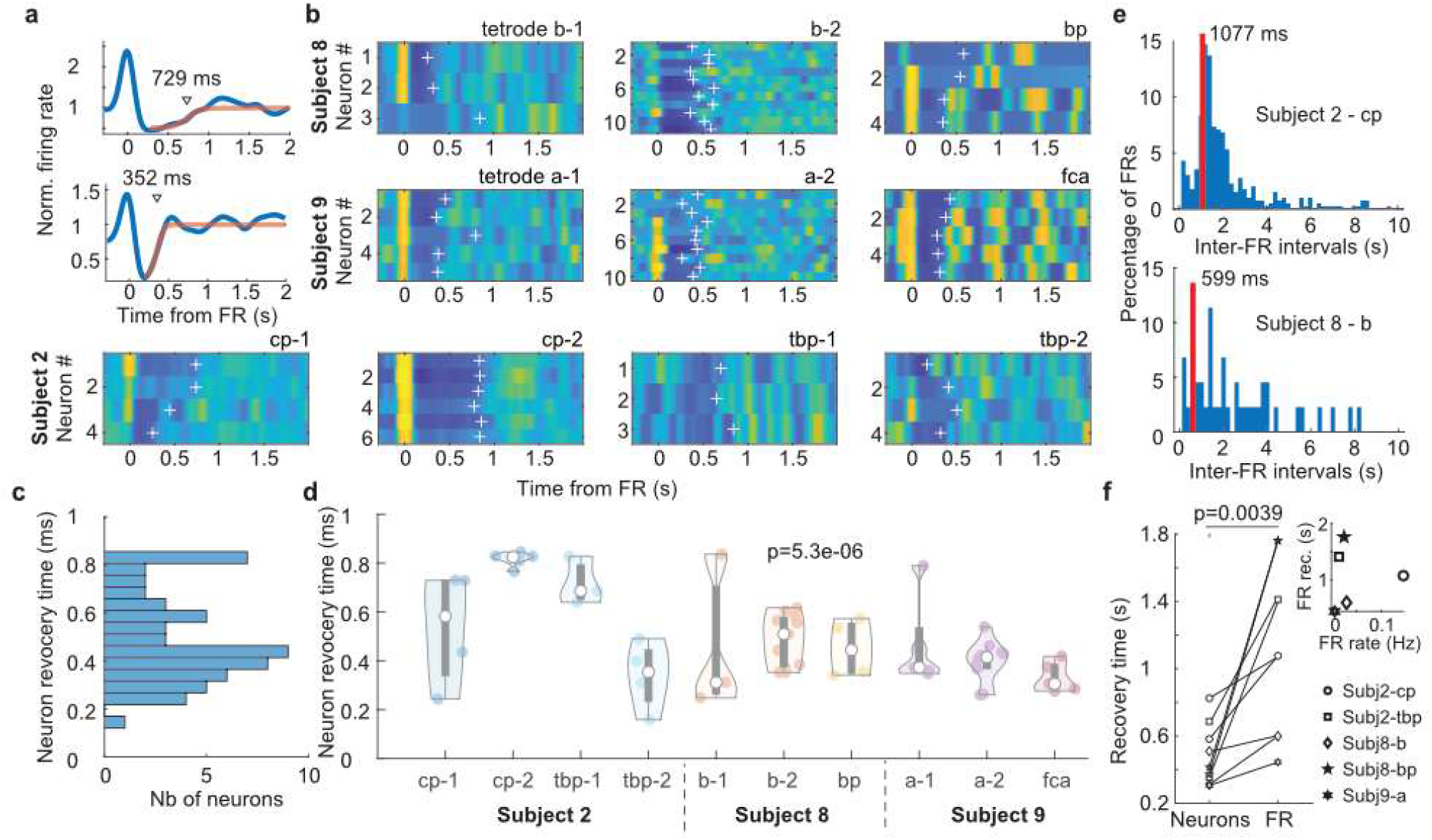
Post-FR recovery of local network states. **a**. Identification of post-FR neuronal recovery time by fitting post-FR neuronal cross-correlograms with sigmoids. Cross-correlograms are normalized to the neuron’s session-wide average firing rate. **b**. Tetrode-by-tetrode post-FR neuronal cross-correlograms in three subjects (total of six hybrid electrodes and 55 units with a significant sigmoid fit in the range 50-1500ms). White crosses indicate recovery time (obtained as in **a**). **c**. Distribution of neuronal recovery time in three subjects. **d**. Distribution of tetrode-by-tetrode recovery times (same presentation as in **Fig. 4b**). **e**. Inter-FR interval distribution from two hybrid electrodes (two subjects). **f**. Hybrid-by-hybrid neuronal and FR recovery times. *Inset*, FR recovery time as a function of FR occurrence rate (r = 0.06; p = 0.9; Pearson’s correlation).

To unravel a potential relationship between neuronal and network-level dynamics, we focused our analysis on three subjects with at least four neurons on two different tetrodes from the same hybrid and showing FRs on these tetrodes (**Fig. 5b**). While post-FR recovery times were broadly distributed across all three recordings, from 282 ms (10^th^ percentile) to 816 ms (90^th^ percentile) (**Fig. 5c**), there was less variability within a tetrode than across tetrodes (**Fig. 5d**; p = 5 × 10^−6^; Kruskal-Wallis test), sometimes even on the same hybrid (see, for example, *cp* and *tbp* hybrids in *subject 2*). Next, we determined the typical recovery time of FRs as the median of all inter-FR intervals shorter than 10 s (**Fig. 5e**). We then determined the circuit-specific neuronal recovery time as the median of recovery times of all neurons recorded on a given tetrode. Neuronal recovery times were of the same order of magnitude as FR recovery time and yet neuronal recovery always preceded FR recovery on a tetrode-by-tetrode basis (**Fig. 5f**; p = 0.0039, n = 9 tetrodes; Wilcoxon signed rank test). These findings suggest that post-FR neuronal inhibition is governed by circuit-specific properties and play an important role in delaying the occurrence of the next FR.

## Discussion

Unraveling the neuronal basis of epileptiform activity is a crucial step towards the development of new and potentially patient-specific diagnostic and therapeutic strategies. To this end, accurate recordings of multiple single units at various anatomical scales is crucial. Decades of animal research have pointed to the advantages of polytrode recordings, especially with tetrodes, to separate extracellular waveforms of multiple single units ^37,42^. Here, by using new hybrid electrodes containing 2-3 tetrodes implanted intracranially in epileptic patients, we characterized some fundamental aspects of neuronal dynamics at times of interictal discharge (**Fig. 2**). Specifically, we demonstrated that FRs, the most critical phase of IEDs, were associated with increased neuronal activity in the local network but not in remote anatomical locations (**Fig. 3**). In contrast, FRs were followed by neuronal inhibition in brain-wide networks. Neuronal modulation by FRs were homogeneous in local networks but differed across hybrid electrodes (i.e. brain areas), even in the same subjects (**Fig. 4**). Finally, FRs were followed by period of decreased firing that were circuit-specific and were tightly related to the inter-FR intervals (**Fig. 5**). These findings shed light on the neuronal mechanism underlying interictal activity generation, as well as on the mechanism preventing the propagation of epileptiform activity and thus possibly delaying the progression of the disease.

The present study could not address some key issues regarding neuronal and epileptiform activity in subjects. First, the recordings were limited in time and could only focus on interictal discharges. How the neuronal dynamics at times of FRs are related to ictal discharges remains unknown. Next, the number of subjects included in the current study was too limited to explore the precise link between neuronal dynamics around the time of FRs and the anatomy, that is, assessing potential differences in neuronal firing in different brain areas. Last, the total number of neurons was still too limited to explore potential differences in cell types. However, our data show how neuronal population is organized around FRs across multiple scales, from the local network (<1 mm) to brain-wide neuronal interaction.

Our findings point to a local generation of FRs, with the typical radius of FR generators being 1-2 mm. Specifically, FRs do not always occur concomitantly on all tetrodes of the same hybrid electrodes. Even if they did, they were not necessarily in phase (**Fig. 1**). Furthermore, the transient and synchronous increase in neuronal firing at the time of FRs in the local circuits suggests that FRs are generated by the firing of neuronal ensembles in the vicinity of the electrode showing FRs (**Fig. 2-4**), as is the case for physiological ripples in rodents ^43^. The level of excitation at the time of FRs can strongly vary across the tetrodes of a single hybrid electrode (**Fig. 4**). This observation may result from differences in recording location, e.g. cortical layers, and additional recordings will be necessary to precisely understand the origin of this difference. At any rate, the proposed size of FR generators is in agreement with human anatomical data (e.g. the typical size of cortical neuronal processes ^44^) and the spatial extent of neuronal pairwise coordination in the neocortex ^45^.

The signals recorded on macro-contacts represent the summation of LFPs in a much larger volume than tetrode microwires. The presence of FRs on macro-contacts can thus mean two things: that a large portion of tissue around the macro-contact generates FRs with random phases or that a smaller volume generates FRs that are precisely in phase ^46^. In general, however, fewer FRs are detected with macroelectrodes than microelectrodes: the ratio of FRs recorded in the seizure-onset zone to the FRs in non-seizure onset zones is higher for microelectrodes than for macroelectrodes ^25^. In some cases, FRs can be recorded exclusively in the EZ by microelectrodes (especially tetrodes), while none can be detected on the macroelectrodes ^38^. These observations are in line with our findings that FR generators are certainly of limited anatomical size. LFPs recorded on microwires thus provide key insights into the dynamics of the local tissue, potentially informing on the origin of the disorder ^47^.

The transient increase in neuronal firing at times of FRs in local networks is followed by a suppression of neuronal activity (**Fig. 3,5**). This is likely the result of inhibitory neuron recruitment by the FRs, as observed in rodents *in vitro* ^48^ and *in vivo* after physiological ripples ^27,29^ and FRs ^49^. Similar observations were obtained from human resected tissue ^50^. The suppression of neuronal activity after FRs was slow (200-800 ms, **Fig. 5**), suggesting that this effect is mediated, at least in part, by slow GABA receptors, such as GABAB ^27,48,51^. The range of these time constants are in agreement with *in vitro* studies ^48,50^. The inter-tetrode variance of the neuronal recovery duration can result from several factors, such as different synaptic dynamics and neuronal excitability ^48^, but the precise mechanism remains to be elucidated.

Functionally, the decrease in neuronal activity following FRs delays the occurrence of the next FR when the network is particularly excitable, playing the role of a refractory period for the network ^27,48,52–54^. Indeed, the typical minimal duration between two consecutive FRs was always longer that the duration of activity suppression, and yet the timescales of neuronal spiking and FR recovery (i.e. time to reach baseline occurrence rate) were of the same order of magnitude and specific to each local network – even within the same subject (**Fig. 5**) – demonstrating a link between fine grained neuronal dynamics and network-level events. Neuronal and FR recovery rate were not significantly correlated per se, in part due to the limited number of observations. However, this also supports the idea that recovery and neuronal activity is only one of several processes leading to the generation of another population burst ^48^. Inhibition-induced refractoriness in local networks has been suggested to play a critical role in the propagation of interictal discharge across areas ^55^ and future work will unravel this phenomenon at the neuronal level.

Interictal epileptic discharges can have highly variable spatiotemporal profiles ^56^, as seen for example on two neighboring tetrodes in **Fig. 2a**. Here, we have shown that neuronal response to FRs was network-specific in terms of excitation level, overall temporal profile of neuronal modulation, and post-FR firing inhibition, confirming earlier reports of high variability in neuronal responses to IEDs ^34,35^. However, different mechanisms may underlie the generation of IEDs and FRs ^57^. Further investigation of the precise relationship between local neuronal activity and associated events in the LFP (IED with or without FRs) will be necessary to guide future clinical investigation of epileptic tissue with IED and IED-concurrent FRs.

The multi-hybrid electrode implants allowed us to address the question of the long-range coordination of neuronal activity around the time of FRs. The strong excitation associated with FRs must have consequences in networks beyond the local circuits where FRs are generated ^58^, as seen, for example, in the cortex after hippocampal ripples in rodents ^59–61^. However, in general, neurons in other brains areas do not show increased excitation at the time of FRs (**Fig. 3**). In contrast, neuronal activity tends to be suppressed globally after FRs (**Fig. 3**). Although the level of activity suppression is lower than what is observed locally (i.e. after FRs in the local circuit), this observation begs the question of the underlying anatomical pathway mediating this long-range activity suppression. One possibility is that this effect is mediated by feedforward inhibition akin to the strong recruitment of fast-spiking inhibitory neurons after hippocampal ripples in rodents ^62^. Importantly, feedforward inhibition has been suggested to be a key mechanism controlling the spread of ictal wavefront ^63–65^. Another possibility is that post-FR activity suppression is mediated by long-range inhibitory neurons ^66,67^. Long-range activity suppression may also result from an indirect network effect, for example, through cortico-thalamo-cortical pathways. This is supported by the triggering of specific thalamocortical oscillations by physiological ripples ^62,68^ and by FRs in rodents and human subjects ^69^. Further work will be necessary to investigate the relationship between FR-triggered long-range activity suppression and the precise anatomical pathway of generalized ictal activity.

In conclusion, the new hybrid electrodes used in the present study offer a unique access to the neuronal dynamics underlying epileptiform activity. Tetrodes enable the characterization of neuronal activity in micro- and macro-circuits. By detecting more local FR generators, combining tetrodes to macroelectrodes in clinical routine may define more accurately the extent of EZ and improve the trust in the outcome of neurosurgery.

## Materials and Methods

### Surgery

The epileptic patients underwent stereoelectroencephalography (SEEG) ^70^ in the Neurophysiological Exploration unit in Toulouse, between March 2015 and May 2019. They all accepted to participate in a prospective protocol (EpiFar project - ClinicalTrials.gov identifier: NCT02491476) aiming to assess the clinical interest of fast ripples using a new type of hybrid micro-macro intracranial electrode (Dixi Medical, Besançon, France) ^36,38^. Every patient suffered from focal epilepsy and antiseizure medication failed to control their seizures. The exact location of the EZ could not be specified by non-invasive assessments in all patients (including MRI, video-EEG, functional imagery such as 18-FDG positron emission tomography in every patient, and additional ictal single positron emission cerebral tomography in some patients). Lesions and the etiologies of epilepsies were heterogeneous across patients (see Supplementary Table 1). SEEG recordings were performed to accurately define the EZ and carried out as part of the patients’ clinical care. The antiseizure medication was gradually reduced to facilitate seizure occurrence in the several days following each implantation, according to the usual clinical procedure. Each subject received detailed information about the objectives of the SEEG technique before intracerebral electrode implantation and about the use of hybrid electrodes. They signed an informed consent form for the implantation and use of the EEG data for research purposes. Implantation of the hybrid electrodes (maximum of 4 per subject) and analyses of intracranial EEG data were approved by the local ethics committee and French Drug and Health Products Safety Agency (CPP Sud-Ouest et Outre-Mer I ethical board, no. 1-14-23, and ANSM 2014-A00747-40).

### Electrodes

SEEG recordings were performed using intracerebral multiple-contact depth electrodes implanted intracranially according to Talairach’s stereotactic method ^71^. The location of each electrode contact was based on a post-operative CT scan/pre-operative 3D T1-weighted MRI data co-registration. The resolution of these data fusions allows a visual control of the anatomical location of each contact and its location in the gray or white matter. The choice of electrode location was based solely on pre-SEEG clinical observations and on hypotheses about the location of the EZ based on non-invasive assessment. The global electrode plan was specific for each patient and combined semi-rigid multi-lead clinical depth macroelectrodes (Microdeep, DIXI Medical, France) and hybrids electrodes. The macroelectrodes had a diameter of 0.8 mm and contained 5-18 contacts (platinum/iridium) 2 mm long and 1.5 mm apart. Two to four hybrid electrodes were implanted by subjects depending on clinical and anatomical constraints (e.g. intra-parenchymal length vs. available length of electrodes). The hybrid electrodes were developed by Dixi Medical, Besançon, France. Implants in human subjects were authorized by the local ethics committee and French Drug and Health Products Safety Agency (CPP Sud-Ouest et Outre-Mer I ethical board, no. 1-14-23, and ANSM 2014-A00747-40). These electrodes were previously described ^38^. Briefly, they are similar in design to regular depth electrodes (diameter: 0.8 mm, length: 33.2, 40.4, or 50.8 mms, depending on the configuration, 6-9 macro-contacts) and equipped with two or three tetrodes (four microelectrodes each) that protruded 3 mm from the shaft between the first and second most medial macro-contacts. Each tetrode is made by twisting four microwires (tungsten, 20 µm) and connected to a specific connector. The tetrodes are extruded by rotating an external micrometer screw immediately after implantation in the operatory room. In this study, tetrodes were all located between the first two macro-contacts (from the tip of the depth electrode).

### Recording system

Macro-contact signals were recorded using two SystemPLUS EVOLUTION 64-channel acquisition units (Micromed, Treviso, Italy) at a sampling rate of 2048 Hz (anti-aliasing filter: 926.7 Hz; high-pass filter: 0.15 Hz; low-pass filter: 1000 Hz). We used the raw SystemPLUS signal to avoid unnecessary filtering ^72^. Simultaneously, microelectrode signals were continuously acquired using a 64-channel Cerebus System amplifier (Blackrock Microsystems, Salt Lake City, UT, USA) at a sampling rate of 30 kHz (0.3-7.5 kHz bandwidth) and time synchronized with the macro-contact signal. Line noise cancellation at 50 Hz was applied. A macro-contact located in the white matter was chosen as a reference for both systems. Macro-contacts were recorded 24/24 hours. Microelectrodes were recorded for 1 hour in five morning sessions, while the subject watched TV.

### Spike sorting

Raw electrophysiological recordings were preprocessed within the Brainstorm environment ^73^. Semi-automatic spike sorting was performed offline on each separate tetrode using SpyKING CIRCUS with a template matching-based algorithm ^74^ on a one-hour signal. Different quality metrics were considered: interspike interval, scatterplots of the different clusters, amplitude over time, auto-correlogram, cross-correlogram, and refractory period violations to optimize spike sorting ^75^. An action potential was detected if the amplitude was above a threshold set at six times the median of the absolute deviation of the voltage. The resulting data were followed by manual curation of the data in Klusters software ^76^.

### Ripple/Fast-ripple detection

Thirty-four subjects were included in the EpiFar project during the inclusion period. A preselection of EEG data for further coupled fast-ripples and neuronal analyses was based both on the results of the spike sorting (see below) and on the results of the FR detection on 10-minute samples on day 3 and 4 after implantation in these 34 subjects. The selection of subjects in the current research was based on (1) the quality of EEG signal that may be unanalyzable in some cases due to a low signal-to-noise ratio, (2) fast ripple visualization on a 10-minute signal sample during preselection, and (3) visible neuronal activity with the possibility of clustering. Depending on these criteria, we selected one hour of recording in 9 subjects, which consists in 26 hybrids for further neuronal and fast-ripples analyses.

Visual analysis of high frequency oscillations is still considered the gold standard for their assessment ^19,77^. However, visual analysis might be subjective and is highly time consuming. On the contrary, automatic detectors have highly heterogeneous performances and none have been validated on human tetrode recordings. Thus, we decided to combine automatic detection of FRs and visual analysis ^38^. FRs were visually and automatically analyzed during 10-minute epochs in each subject during the 3rd day and the 4th day of recordings during the waking period. This duration of sample recordings is at least similar or even longer than previous publications ^19,77^. We voluntarily chose a duration of analysis across 2 days since the FR rate can be highly variable across days, especially in relation to the progressive withdrawal of treatment. At the 3rd and 4th days, subjects’ medication was in most cases reduced. In parallel, this FR detection for preselection was training to better isolate FRs manually on a longer signal duration for further analyses. There were three criteria for identifying FRs: (1) an oscillation visible in both the raw data and the filtered signal (200-600 Hz bandpass finite impulse response filter to reduce ringing); (2) at least four oscillations with an amplitude clearly higher than the baseline ^78^; and (3) time-frequency analysis resulting in circumscribed areas in the time-frequency plane in the FR band to avoid artifacts and false ripples created by filtering ^72,79^. Monopolar montages and reformatted bipolar montages were used for micro- and macroelectrode recordings, respectively. First, we used the automatic detector Delphos ^80^ within the Anywave software ^81^ on all 10-minute epochs. It is an automatic detector that improves the detectability of HFO by applying a linear whitening (i.e. flattening) transformation to enhance the fast oscillations while preserving an optimal signal-to-noise ratio. Then, each event detected by Delphos was manually reviewed to verify that all previous criteria were met. The raw and filtered signals (200-600 Hz) of the four microwires of each tetrode were visualized simultaneously using a 0.6 s time window. FRs were detected independently by two researchers (JC, ED) blind to clinical conclusions and healthy regions, as well as to the final decision regarding neurosurgery at the moment of fast ripple detection. Events without the 3 previous criteria were systematically rejected. Noisy periods or channels were excluded. Then, in the 9 selected subjects, FRs were manually tracked in Brainstorm ^82^on a one-hour signal on the microelectrodes (see criteria for identification below).

### Determination of the epileptic network

In clinical practice, the epileptogenic zone (EZ) is an electroclinical definition corresponding to the networks of brain regions generating seizures ^83,84^. The complete removal of the EZ is essential to prevent seizures after surgery ^1^. EZ is supposed to integrate the seizure-onset zone. The irritative zone (IZ) is the site of IED ^3,13,85^. The propagation zone refers to brain areas where secondary delayed electrophysiological discharges are observed outside the EZ. Healthy brain tissue (O) are the areas with no overlap of EZ and IZ and where no seizure propagates. Determination of the clinical EZ and non-EZ as well as the IZ, propagation zone, and healthy areas was made independently by the clinical team of epileptologists after reviewing the clinical iEEG data (LV, MD). They were blind to FRs and neuronal analyses. All the neuronal and FR analyses were primarily done by JC and AP blind to final clinical conclusions.

### Motion tracking

Tracking of each subject’s movement during recording was obtained from a single video camera (at 25 Hz) whose acquisition was synchronized with electrophysiological data (Micromed, Treviso, Italy) in three subjects (1, 6, and 8). Videos were processed post-hoc using Deeplabcut ^39^ to analyze subjects’ movements (**Fig. 1**). In each subject, 6 points were tracked: nose, chin, left/right shoulders, and left/right hands. A Resnet50 network was trained for at least 100,000 iterations or longer if tracking was not satisfactory. The velocity of each marker was computed and smoothed with a 1-second Gaussian kernel. The velocity for time stamps at which the log-likelihood of marker detection was below 0.95 was considered null. Overall movement velocity was computed as the average velocity of all markers. Subjects were considered still if the average marker velocity was below 25 pixels/second for at least 3 seconds.

### Computing z-scored cross-correlograms

Cross-correlograms reflect the average firing rate of a neuron relative to the timing of FRs, and are thus biased by the neuron’s firing rates. To normalize fluctuation and compare neurons of different firing rates, cross-correlograms were z-scored following a two-step procedure, as described by Viejo and Peyrache (2020).

### Comparison of z-scored cross-correlograms (Fig. 4c)

To evaluate similarity and differences in temporal profiles of neuronal modulation around the time of FRs, we first transformed the cross-correlograms with a non-linear function (*Arctan*) so that the main effect is not dominated by differences in absolute activation. We then projected the transformed cross-correlograms on the first two principal components of the cross-correlograms (accounting for ∼90% of the total variance). Finally, for each neuron, we computed their Mahalanobis distance to all other neurons of the same tetrode and of the other tetrode of the same hybrid electrode, as well as or the mean distance to tetrodes of other hybrid electrodes of the same subject.

## Acknowledgments

We thank Karine Bouyer, Simona Celebrini, Pascale Cook, Martin Deudon, Ludovic Gardy, Leila Reddy, Simon J. Thorpe, for helpful comments and help with data acquisition; Sara Mahallati, Adrian Duszkiewicz, and Karim Benchenane for discussion and comments on the manuscript. This project has been made possible with the financial support of the European Research Council (under the European Union’s Seventh Framework Programme (FP/2007-2013) / ERC Grant Agreement no. 323711, M4 project), the Fondation Française pour la Recherche sur l’Epilepsie (Prix Marion Clignet), Toulouse University Hospital (through a call for technological and organisation innovation research projects), and Health Canada (through the Canada Brain Research Fund), an innovative partnership between the government of Canada (through Health Canada), and Brain Canada (to AP). This work was also supported by a CIHR Project Grant (155957), NSERC Discovery Grant (RGPIN-2018-04600), and a Canada-Israel Health Research Initiative, jointly funded by the Canadian Institutes of Health Research, the Israel Science Foundation, the International Development Research Centre, Canada and the Azrieli Foundation (108877-001) to AP, as well as a fellowship from the Ligue Française Contre l’Epilepsie to JC.

