## Supplementary Figure 1 and Table 1 for "Local neuronal excitation and global inhibition during epileptic fast ripples in humans" (from another version of this preprint: version 1, 2021-09-11)

### Supplementary Information.

Supplementary Figure 1 and Supplementary Table 1.

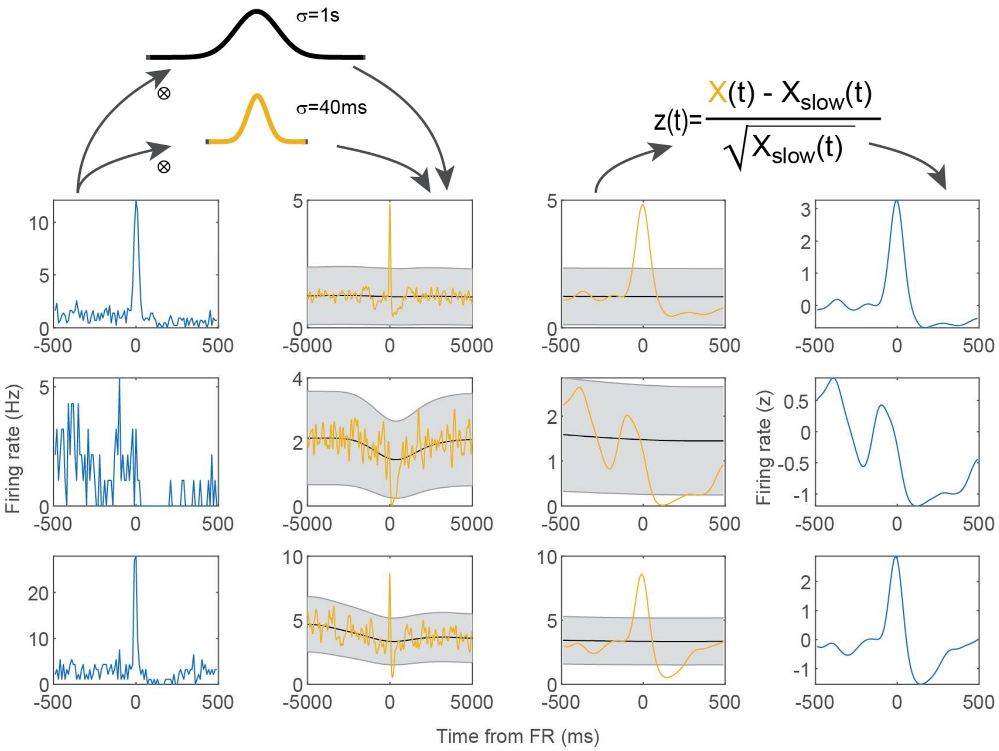


**Supplementary Fig. 1 | Cross-correlogram normalization and z-scoring**. The raw cross-correlogram of neuronal activity in relation to FRs (1st column) is low-pass filtered in two different ways (2nd and 3rd columns): with a short Gaussian kernel (SD 40 ms, yellow curves) to reduce bin-to-bin variability and with a long Gaussian kernel (SD 1 s, black curve) to estimate the instantaneous baseline firing rate. Assuming a Poisson process, the baseline SD is taken as the square root of the baseline firing rate. The normalized cross-correlograms were obtained by subtracting the baseline firing rate from the short timescale low-pass filtered signal and divided by the square root of the baseline firing rate, yielding z-scored firing rates. Note that 3rd column is the same as the 2nd one but at the same time scale as the 1st and 4th columns.

| Subject | Hybrid name | Tetrodes | Neurons (number) | Fast-ripples (number) | Anatomical location | Location related to epileptic network | Lesion type | Gender | Age |
|---|---|---|---|---|---|---|---|---|---|
| 1 | a | a1 | 0 | 0 | Amygdala, R | IZ | Hippocampal sclerosis | M | 41 |
| | | a2 | 3 | | | | | | |
| | | a3 | 3 | | | | | | |
| | bp | bp1 | 0 | 139 | Anterior hippocampus, L | EZ | | | |
| | | bp2 | 0 | | | | | | |
| | ap | ap1 | 7 | 129 | Amygdala, L | EZ | | | |
| | | ap2 | 2 | | | | | | |
| | | ap3 | 2 | | | | | | |
| | tbp | tbp1 | 0 | 281 | Rhinal cortex, L | EZ | | | |
| | | tbp2 | 0 | | | | | | |
| 2 | tb | tb1 | 0 | 0 | Rhinal cortex, R | IZ | Encephalitis sequelae | M | 55 |
| | | tb2 | 0 | | | | | | |
| | cp | cp1 | 4 | 430 | Posterior hippocampus, L | EZ | | | |
| | | cp2 | 7 | | | | | | |
| | | cp3 | 0 | | | | | | |
| | ap | ap1 | 0 | 74 | Amygdala, L | EZ | | | |
| | | ap1 | 0 | | | | | | |
| | tbp | tbp1 | 3 | 42 | Rhinal cortex, L | EZ | | | |
| | | tbp2 | 9 | | | | | | |
| 3 | a | a1 | 0 | 13 | Amygdala, R | IZ | Hippocampal sclerosis | M | 44 |
| | | a2 | 0 | | | | | | |
| | tpp | tpp1 | 0 | 0 | Temporal pole, L | EZ | | | |
| | | tpp2 | 0 | | | | | | |
| | ap | ap1 | 0 | 24 | Amygdala, L | IZ | | | |
| | | ap2 | 0 | | | | | | |
| | cp | cp1 | 1 | 4 | Posterior hippocampus, L | IZ | | | |
| | | cp2 | 1 | | | | | | |
| 4 | gl | gl1 | 2 | 0 | Lingual gyrus, R | O | Cortical dysplasia | F | 36 |
| | | gl2 | 3 | | | | | | |
| | glp | glp1 | 0 | 0 | Lingual gyrus, L | O | | | |
| | | glp2 | 4 | | | | | | |
| | otp | otp1 | 0 | 0 | Fusiform gyrus, L | ZI | | | |
| | | otp2 | 0 | | | | | | |
| | tbp | tbp1 | 0 | 23 | Rhinal cortex, L | ZE | | | |
| | | tbp2 | 3 | | | | | | |
| | | tbp3 | 0 | | | | | | |
| 5 | h | h1 | 0 | 0 | Superior temporal gyrus, R | ZI | Hippocampal sclerosis, cortical dysplasia | M | 16 |
| | | h2 | 8 | | | | | | |
| | b | b1 | 0 | 30 | Anterior hippocampus, R | ZE | | | |
| | | b2 | 0 | | | | | | |
| | ot | ot1 | 0 | 0 | Fusiform gyrus, R | ZI | | | |
| | | ot2 | 0 | | | | | | |
| | cp | cp1 | 0 | 0 | Posterior hippocampus, L | O | | | |
| | | cp2 | 0 | | | | | | |
| 6 | tbp | tbp1 | 0 | 4 | Amygdala, L | IZ | Cortical development abnormality | M | 58 |
| | | tbp2 | 0 | | | | | | |
| | b | b1 | 0 | 0 | Anterior hippocampus, R | O | | | |
| | | b2 | 0 | | | | | | |
| | bp | bp1 | 0 | 286 | Anterior hippocampus, L | IZ | | | |
| | | bp2 | 0 | | | | | | |
| | tpp | tpp1 | 0 | 0 | Temporal pole, L | ZI | | | |
| | | tpp2 | 0 | | | | | | |

| | | | | | | | | | |
|---|---|---|---|---|---|---|---|---|---|
| 7 | tb | tb1 | 0 | 0 | Rhinal cortex, R | IZ | Meningo-encephalocele | M | 54 |
| | | tb2 | 8 | | | | | | |
| | tpp | tpp1 | 15 | 0 | Temporal pole, L | EZ | | | |
| | | tpp2 | 2 | | | | | | |
| | cp | cp1 | 0 | 26 | Posterior hippocampus, L | EZ | | | |
| | | cp2 | 0 | | | | | | |
| | | cp3 | 0 | | | | | | |
| | tbp | tbp1 | 0 | 0 | Rhinal cortex, L | EZ | | | |
| | | tbp2 | 1 | | | | | | |
| 8 | b | b1 | 0 | 93 | Anterior hippocampus, R | EZ | Hippocampal sclerosis | F | 49 |
| | | b2 | 4 | | | | | | |
| | | b3 | 12 | | | | | | |
| | c | c1 | 0 | 63 | Posterior hippocampus, R | EZ | | | |
| | | c2 | 0 | | | | | | |
| | | c3 | 4 | | | | | | |
| | bp | bp1 | 0 | 13 | Anterior hippocampus, L | IZ | | | |
| | | bp2 | 4 | | | | | | |
| 9 | fca | fca1 | 5 | 18 | Posterior medial temporal, R (within nodular heterotopia) | EZ | Nodular heterotopia | M | 26 |
| | | fca2 | 0 | | | | | | |
| | a | a1 | 5 | 71 | Amygdala, R | IZ | | | |
| | | a2 | 0 | | | | | | |
| | | a3 | 9 | | | | | | |
| | tb | tb1 | 0 | 0 | Rhinal cortex, L | IZ | | | |
| | | tb2 | 0 | | | | | | |
| | ap | ap1 | 4 | 0 | Amygdala, L | IZ | | | |
| | | ap2 | 2 | | | | | | |

**Supplementary Table | Patients' clinical characteristics, location of hybrids, and yield of neurons and FRs per tetrode**. *IZ: irritative zone, EZ: epileptogenic zone, O: healthy tissue, L: left, R: right, F: female, M: male.*
